# Circadian regulation of glutamate release pathways shapes synaptic throughput in the brainstem nucleus of the solitary tract (NTS)

**DOI:** 10.1101/2022.10.28.514250

**Authors:** Forrest J. Ragozzino, BreeAnne Peterson, Ilia N. Karatsoreos, James H. Peters

## Abstract

Circadian regulation of autonomic reflex pathways pairs physiological function with the daily light cycle. The brainstem nucleus of the solitary tract (NTS) is a key candidate for rhythmic control of the autonomic nervous system. Here we investigated circadian regulation of NTS neurotransmission and synaptic throughput using patch-clamp electrophysiology in brainstem slices from mice. We found that spontaneous quantal glutamate release on to NTS neurons showed strong circadian rhythmicity, with the highest rate of release during the light phase and the lowest in the dark, that were sufficient to drive day / night differences in constitutive postsynaptic action potential firing. In contrast, afferent-evoked action potential throughput was enhanced during the dark and diminished in the light. Afferent-driven synchronous release pathways showed a similar decrease in release probability that did not explain the enhanced synaptic throughput during the night. However, analysis of postsynaptic membrane properties revealed diurnal changes in conductance; which, when coupled with the circadian changes in glutamate release pathways, tuned synaptic throughput between the light and dark phases. These coordinated pre- / postsynaptic changes encode nuanced control over synaptic performance and pair NTS action potential firing and vagal throughput with time of day.

## INTRODUCTION

Nearly all physiological processes and behaviors are synchronized to the environmental light cycle (Challet, 2019). The suprachiasmatic nucleus (SCN) of the hypothalamus is the master circadian clock in mammals as it receives direct sensory input from the retina, integrating daily changes in ambient light with anticipatory behavioral and physiological changes of the organism (Hastings et al., 2018). SCN rhythmicity is coupled through a cell-autonomous transcription-translation feedback loop (TTFL) consisting of the rhythmic activity of BMAL and CLOCK transcription factors on E-box promotor regions (Takahashi, 2017). The output of this molecular clock controls circadian rhythmicity in many aspects of physiology; including calcium signaling, membrane potential, synaptic transmission, and endocrine signaling (Harvey et al., 2020). SCN generation of these signals is essential for the circadian control of physiological responses, autonomic reflexes, and associated behaviors. Autonomic tone and feeding behavior are strongly under circadian control, and disruption or desynchronization of these physiological rhythms contributing to many disease states including the progressive development of cardiovascular and metabolic dysregulation, as well as the development of obesity (Karatsoreos et al., 2011). While many studies have focused on understanding rhythm generation in the SCN, much less is known about extra-SCN clock mechanisms. The nucleus of the solitary tract (NTS) in the brainstem is a key candidate for the extra-SCN rhythmic control of autonomic regulation, food intake, processes of satiety, and metabolic regulation (Ritter et al., 2017).

The NTS monitors changes in physiological states and coordinates autonomic regulation of bodily homeostasis (Andresen et al., 1994). Information regarding visceral organ status is relayed to the brain via primary vagal afferent neurons which form strong excitatory contacts onto second-order NTS neurons. In addition to neural input, NTS neurons are also responsive to changes in metabolic status and circulating hormones (Appleyard et al., 2005; Baptista et al., 2005; Cui et al., 2012). Outputs of the NTS control many autonomic reflex pathways as well as feeding, satiety and resulting gastrointestinal physiological responses, all of which possess robust circadian oscillations (Buijs et al., 2006; Konturek et al., 2011; Panda, 2016). Previous studies have demonstrated rhythmicity of clock gene expression in the NTS and the cell bodies of vagal afferent neurons (Herichova et al., 2007; Kaneko et al., 2009; Chrobok et al., 2020), suggesting local TTFL clock signaling. However, the cellular mechanisms through which circadian rhythms orchestrate information processing in the NTS remain unclear. We posit circadian nuanced control of synaptic transmission and action potential throughput mediate these daily changes.

Vagal afferent neurons release glutamate via both action potential driven (synchronous and asynchronous) and action potential independent (spontaneous) vesicle release pathways (Wu et al., 2014; Kaeser et al., 2014). While synchronous and asynchronous vesicle release convey real-time action-potential driven signals arising from the viscera, ongoing spontaneous glutamate release sets the synaptic tone and sensitivity to these incoming signals (Shoudai et al., 2010; Kavalali, 2015). It has been well characterized that hormonal and other receptor mediated signals commonly couple to the vesicle release machinery to exert their effects on neural circuits (Appleyard et al., 2005; Bailey et al., 2006; Peters et al., 2008). In this set of studies, we systematically investigate the ability of time of day and circadian regulatory processes to alter pre- and postsynaptic processes to shape constitutive and afferent driven action potential throughput in the NTS.

## RESULTS

### The NTS and nodose ganglia have strongly rhythmic TTFL Clock gene expression (Figure 1)

Tissue was collected from the NTS and both nodose ganglia at six time points across the day (N = 4 - 5 mice / time point, 25 mice total) from animals under 12:12 LD conditions; and samples were processed using RT-qPCR for quantification of TTFL Clock gene expression levels (**Figure 1A**). Expression for *Per1, Per2*, and *NR1D1* (Rev-erbα) showed significant diurnal change in both the NTS and nodose ganglia (**Figure 1B-D**). Whereas *Clock* expression did not significantly change throughout the day (**Figure 1E**), and *Bmal1* (*Arntl1*) was only found to be rhythmic in the nodose tissue (**Figure 1F**).

**FIGURE 1:**
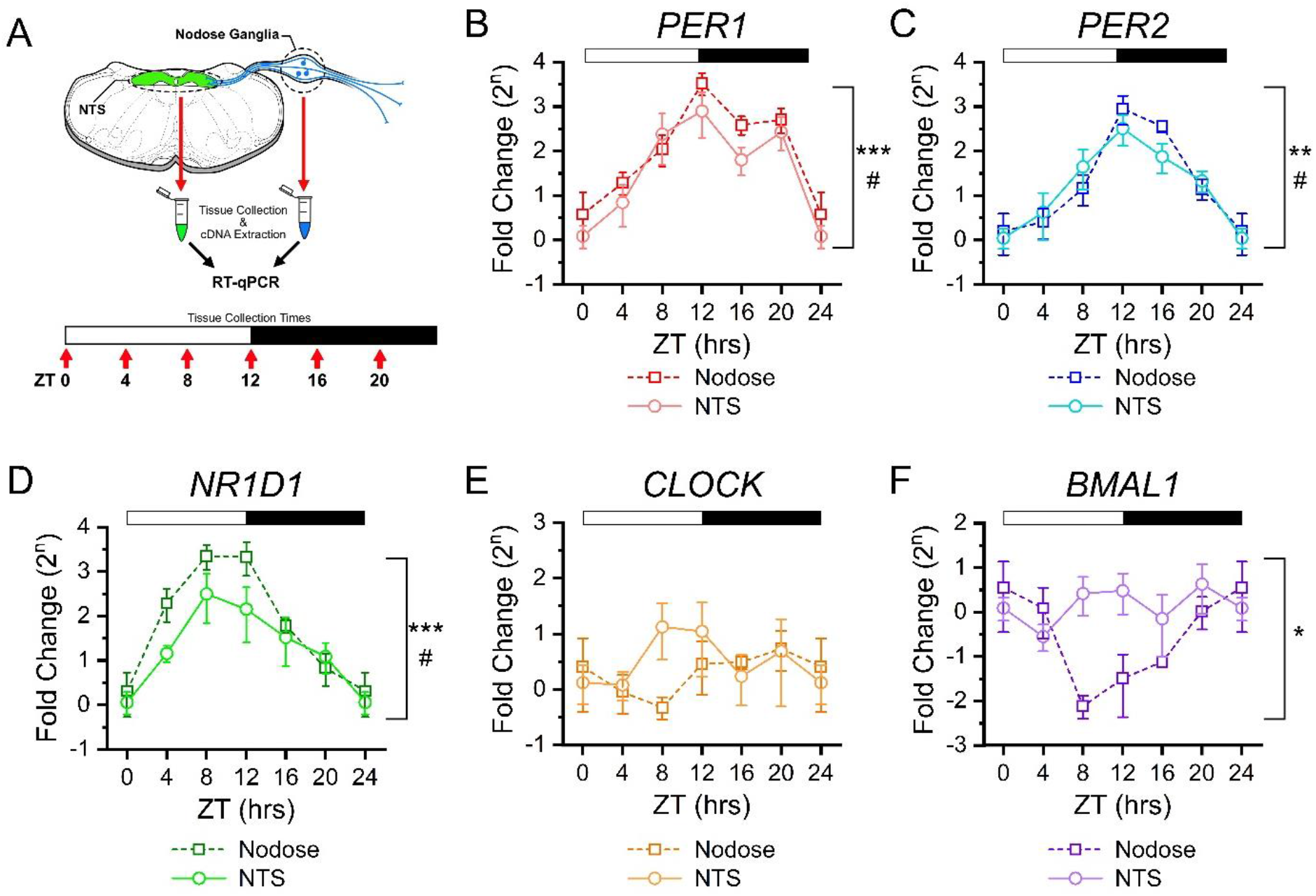
Rhythmic clock gene expression in NTS and vagal afferent neurons across time of day. **A)** Clock gene expression was determined in the NTS and nodose ganglia using RT-qPCR from tissues taken at six time points throughout the light/dark cycle. For each time point tissue was collected from N = 4 - 5 mice and samples were run in triplicate. Canonical clock gene expression from nodose ganglia (squares) and NTS (circles) show rhythmic changes for **B)** *Per1* (NTS: ANOVA, #P = 0.039; Nodose: ANOVA, ***P < 0.001), **C)** *Per2* (NTS: ANOVA, #P = 0.01; Nodose: Kruskal-Wallis, **P = 0.003), **D)** *Nr1d1* (Rev-erbα) (NTS: Kruskal-Wallis, #P = 0.032; Nodose: Kruskal-Wallis, ***P < 0.001); but not for **E)** *Clock* (NTS: ANOVA, P = 0.44; Nodose: ANOVA, P = 0.48), and **F)** only in nodose tissue, but not NTS, for *Arntl1* (Bmal1) (NTS: ANOVA, P = 0.62; Nodose: Kruskal-Wallis, *P = 0.031).^ΔΔ^Ct was calculated relative to RN18s and ZT0, with data expressed as a fold-change on a log base 2 scale.

### Rhythmicity of spontaneous glutamate release onto NTS neurons is under circadian control (Figure 2)

To investigate the circadian regulation of NTS synaptic transmission we first assayed spontaneous glutamate release parameters across the time-of-day using whole-cell voltage-clamp recordings made from NTS neurons in acute *ex vivo* horizontal brainstem slices (**Figure 2A**). We observed marked differences in the frequency of glutamate mediated spontaneous excitatory postsynaptic currents (sEPSCs) between recordings taken during the light-phase (day) compared to the dark-phase (night) (**Figure 2B**). The average frequency of sEPSCs demonstrated diurnal rhythmicity that peaked during late day (ZT6-12) and was at a minimum during late dark phase (ZT18-24; N = 168 neurons / 14 mice across all times points; **Figure 2C, left panel**). Comparison of sEPSCs frequencies in 12 hr bins showed significantly higher rates of release during the day (7.8 ± 0.9 Hz, N = 72 neurons) compared to the night (3.7 ± 0.3 Hz, N = 92 neurons) (**Figure 2C, right panel**). To determine whether this rhythm was under circadian control we replicated this experiment from mice placed in constant dark (DD) conditions for 24-42 hours (one full circadian cycle) prior to time of recording. Consistent with clock control, sEPSC frequency remained higher during the relative day (CT 0-12: 5.5 ± 0.5 Hz, N = 61 neurons) compared to relative night (CT 12-24: 4.0 ± 0.4 Hz, N = 55 neurons) with a similar periodicity (Total N = 116 neurons / 15 mice across all times points) (**Figure 2D**). Waveform analysis revealed sEPSC amplitudes were not statistically different across time of day in either the LD (Day: 29.7 ± 1.4 pA, Night: 30.2 ± 1.4 pA) or DD conditions (Day: 27.4 ± 1.5 pA; Night: 32.0 ± 2.1 pA) (**Figure 2E-F**).

**FIGURE 2:**
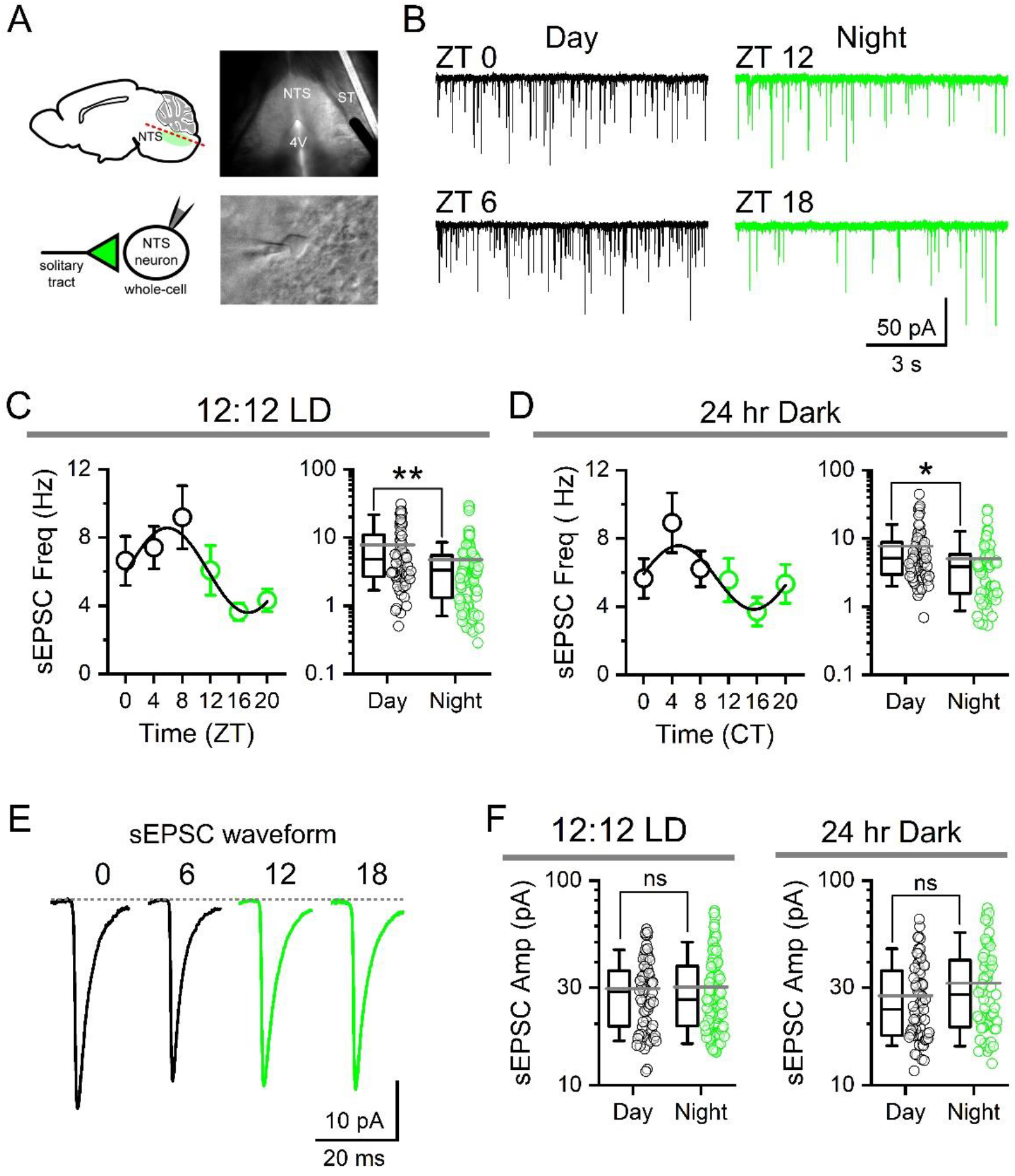
Spontaneous glutamate release frequency shows circadian rhythmicity. **A)** Infrared-DIC images of a mouse horizontal brain slice (top) and whole-cell recording of a single NTS neuron (bottom) during patch-clamp experiments. NTS, nucleus of the solitary tract; ST, solitary tract; 4V, fourth ventricle. Cells were voltage-clamped at −60 mV. **B)** Representative current traces showing spontaneous EPSCs (sEPSCs) from recordings in slices taken at one of four time-points throughout the day: ZT 0, ZT 6, ZT 12, and ZT 18. **C)** Average sEPSC frequency recorded from animals housed under 12:12 LD conditions plotted in four-hour bins across time-of-day showing the change in frequency (left panel). Frequency of release was significantly different during the day (ZT 0-12) when compared to the night (ZT 12-24) (right panel) (N = 72 – 92 neurons / 14 mice, Mann-Whitney Rank Sum Test, **P < 0.001). **D)** Recordings from mice taken after 24-42 hrs of constant darkness (12:12 DD) maintained rhythmicity of sEPSC frequency independent of light cues. Frequency was significantly higher from cells taken during the respective day (CT 0-12) compared to the dark (CT 12-24) (N = 55 – 61 neurons / 15 mice, Mann-Whitney Rank Sum Test, **P = 0.01). **E)** Average sEPSC waveforms across time of day. Traces are averages of 100 discrete events from representative recordings. **F)** There was no significant difference in sEPSC amplitude between day and night in LD (Mann-Whitney Rank Sum Test, P = 0.89) or DD conditions (Mann-Whitney Rank Sum Test, P = 0.14).

To determine whether circadian rhythmicity of sEPSC frequency is secondary to changes in neuronal action potential firing, we replicated our observations in the presence of the voltage activated sodium channel inhibitor tetrodotoxin (TTX, 1 μM) (**Figure 3**). Inhibition of action potential firing produced a modest, but statistically significant, decrease in the frequency of spontaneous glutamate release during both the daytime (ZT 4-9) and nighttime (ZT 16-21) recordings (Total N = 29 neurons / 6 mice) (**Figure 3A-B**). However, TTX exposure did not eliminate the day / night difference in sEPSC frequency (**Figure 3B**). The amplitude of sEPSCs was not changed by TTX, consistent with the quantal nature of spontaneous events in the NTS (**Figure 3C**).

**FIGURE 3:**
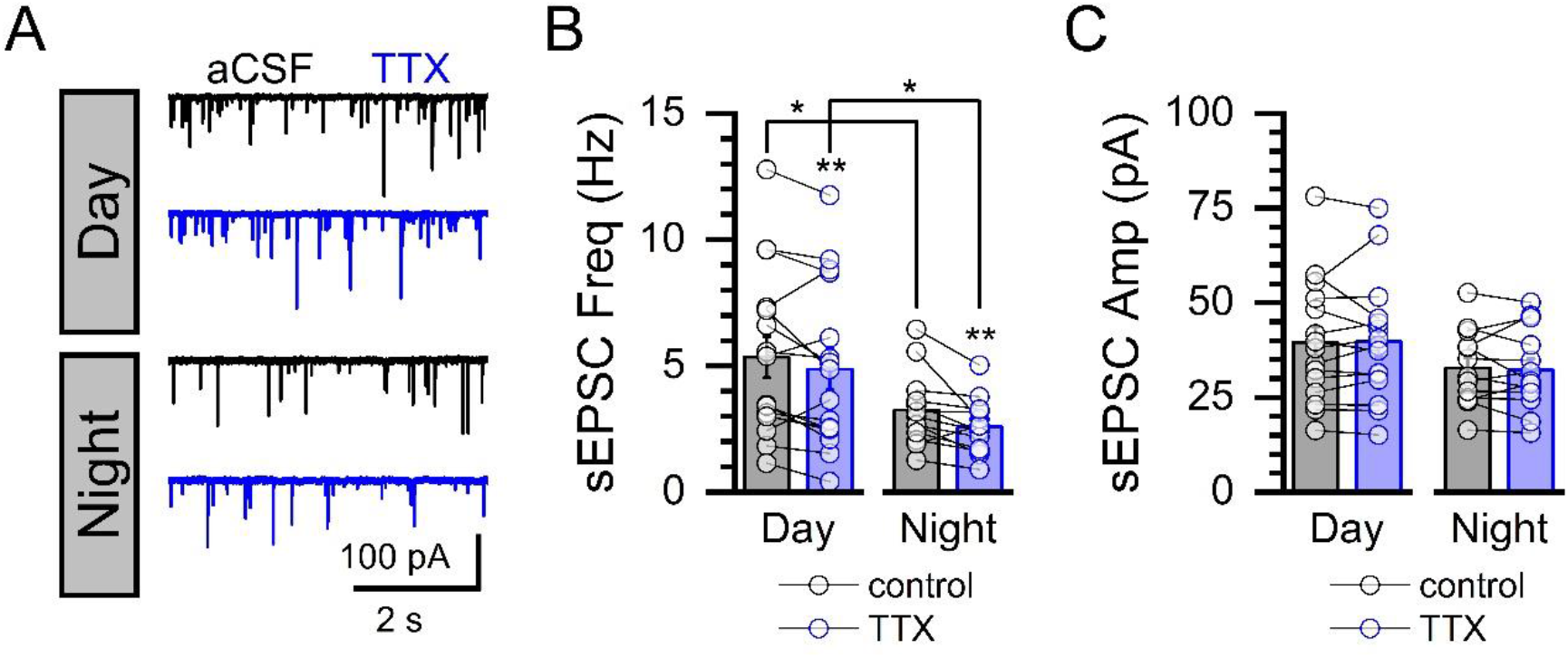
Rhythmic changes in spontaneous glutamate release occur independent of action potential firing. **A)** Representative current traces showing spontaneous events before and after bath perfusion of tetrodotoxin (TTX, 1 μM) during daytime and nighttime recordings. **B)** Bath application of TTX reduced the sEPSC frequency during day (Paired T-test, N = 16 neurons / 3 mice, *P = 0.015) and night (Paired T-test, N = 13 neurons / 3 mice, *P = 0.012) but failed to eliminate the day/night difference (Control Day/Night: Mann-Whitney Rank Sum Test, N = 13 – 16 neurons / 6 mice, *P = 0.04; and TTX Day/Night: Mann-Whitney Rank Sum Test, N = 13 – 16 neurons / 6 mice, *P = 0.05). **C)** Exposure to TTX had no effect on sEPSC amplitude during either the day (Paired T-test, P = 0.80) or night (Paired T-Test, P = 0.35).

### Diurnal variations in NTS action potential firing are driven by glutamate signaling

To determine if the changes in glutamate release were sufficient to drive postsynaptic action potential firing, we next performed cell-attached extracellular and whole-cell recordings in *ex vivo* brainstem slices taken between ZT 4-5 for daytime recordings and ZT 16-17 for nighttime recordings (**Figure 4**). We found the NTS had a greater percentage of neurons spontaneously firing action-potentials at higher frequencies during the day compared to the night (**Figure 4A**). Bath application of ionotropic glutamate receptor antagonists NBQX (25 μM) and D-AP5 (25 μM) eliminated the day / night difference in action potential frequency (**Figure 4A-B**). Further, glutamate receptor antagonism also decreased the proportion of spontaneously firing neurons from 92% to 50% during the day (N = 93 neurons / 6 mice total, Chi-square, ***P < 0.001) and 62% to 33% during the night; eliminating the day / night difference (**Figure 4C**). Whole-cell recordings showed NTS neurons trended toward a more depolarized state and showed higher firing frequencies at rest during the day (**Figure 4D-F**). Together these findings support the role of glutamate to drive rhythmic changes in background NTS neuronal firing; with basal firing greatest during the light and diminished at night.

**FIGURE 4:**
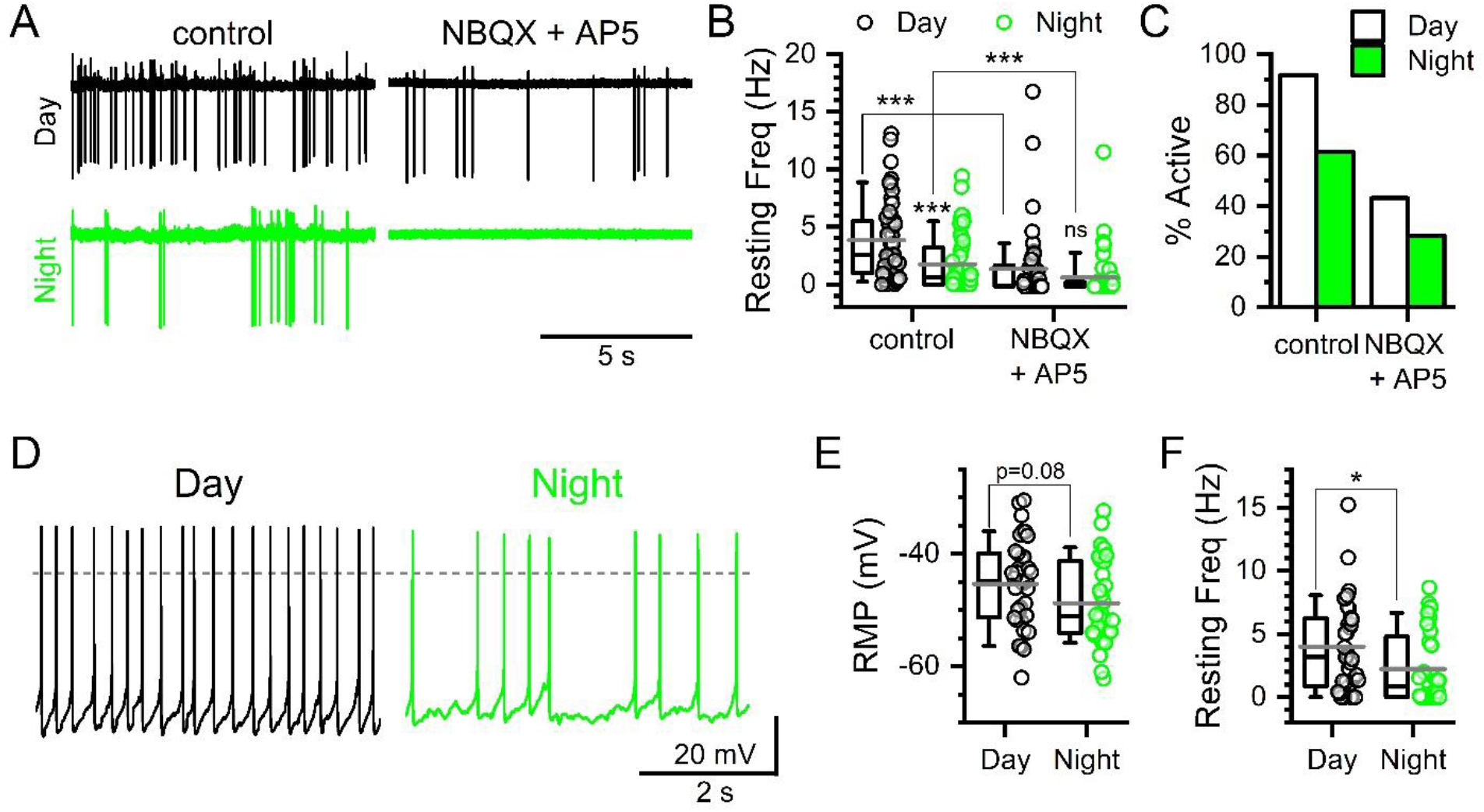
Glutamate drives diurnal rhythmicity in basal action potential firing. **A)** Representative extracellular voltage traces showing spontaneous action potentials during the day (ZT5-9) and night (ZT17-21) under control conditions and following bath application of ionotropic glutamate antagonists (NBQX 25 μM and AP5 25 μM). **B)** Baseline action potential frequency was significantly greater during the day compared to night (Mann-Whitney Rank Sum Test, N = 50 – 53 neurons / 6 mice, ***P < 0.001). Bath application of NBQX + AP5 lowered frequency during the day (Mann-Whitney Rank Sum Test, N = 44 – 50 neurons / 6 mice, ***P < 0.001) and night (Mann-Whitney Rank Sum Test, N = 46 – 53 neurons / 6 mice, ***P < 0.001); eliminating the day / night difference in action potential firing (Mann-Whitney, N = 44 – 46 neurons / 6 mice, P = 0.12). **C)** Similarly, the percent of neurons firing action potentials spontaneously under control conditions was greater during the day compared to the night (Chi-square, N = 103 neurons / 6 mice, ***P < 0.001). This day / night difference was reduced following NBQX + AP5 and no longer statistically significant (Chi-square, N = 100 neurons / 6 mice, P = 0.21). **D)** Whole-cell recordings from NTS neurons confirmed the day / night difference in action potential frequency. **E)** While there was no significant difference in the resting membrane potentials (RMP) (T-test, N = 32 – 35 neurons /, P = 0.08) spontaneous action potential firing frequencies were significantly faster during the day (**F**, Mann-Whitney Ranked Sum Test, N = 32 – 35 neurons /, *P = 0.03).

### Day / night differences in the fidelity of throughput at ST-NTS synapses

The fidelity of synaptic throughput reflects the ‘attention’ of NTS neurons to incoming viscerosensory afferent inputs. Presynaptic activation of ST-afferents produces glutamate dependent action potential firing in NTS neurons (**Figure 5**). The precision of synaptic signaling is the result of both presynaptic release conditions and postsynaptic membrane properties. Similar to previous reports, we found the fidelity of ST-NTS synaptic throughput was high initially, due to elevated initial release probability, but diminished across the stimulus train with a decrease in the precision of action potential initiation and increase in failures becoming more frequent (**Figure 5A**) (Bailey et al., 2006b). Surprisingly, we observed that synaptic throughput was significantly lower in recordings taken during the day compared to the night (**Figure 5B**). To explore this relationship further we repeated the experiment at both higher and lower stimulus intensities. There was no difference in throughput at lower stimulus intensity (0.1 mA), presumably due to be at or below threshold. Nor was there a significant difference at higher intensities (1 or 3 mA), likely due to recruitment of multiple convergent afferent inputs (**Figure 5C**). Stimulation failed to elicit action potentials in the presence of ionotropic glutamate receptor antagonists NBQX and D-AP5 (*data not shown*).

**FIGURE 5:**
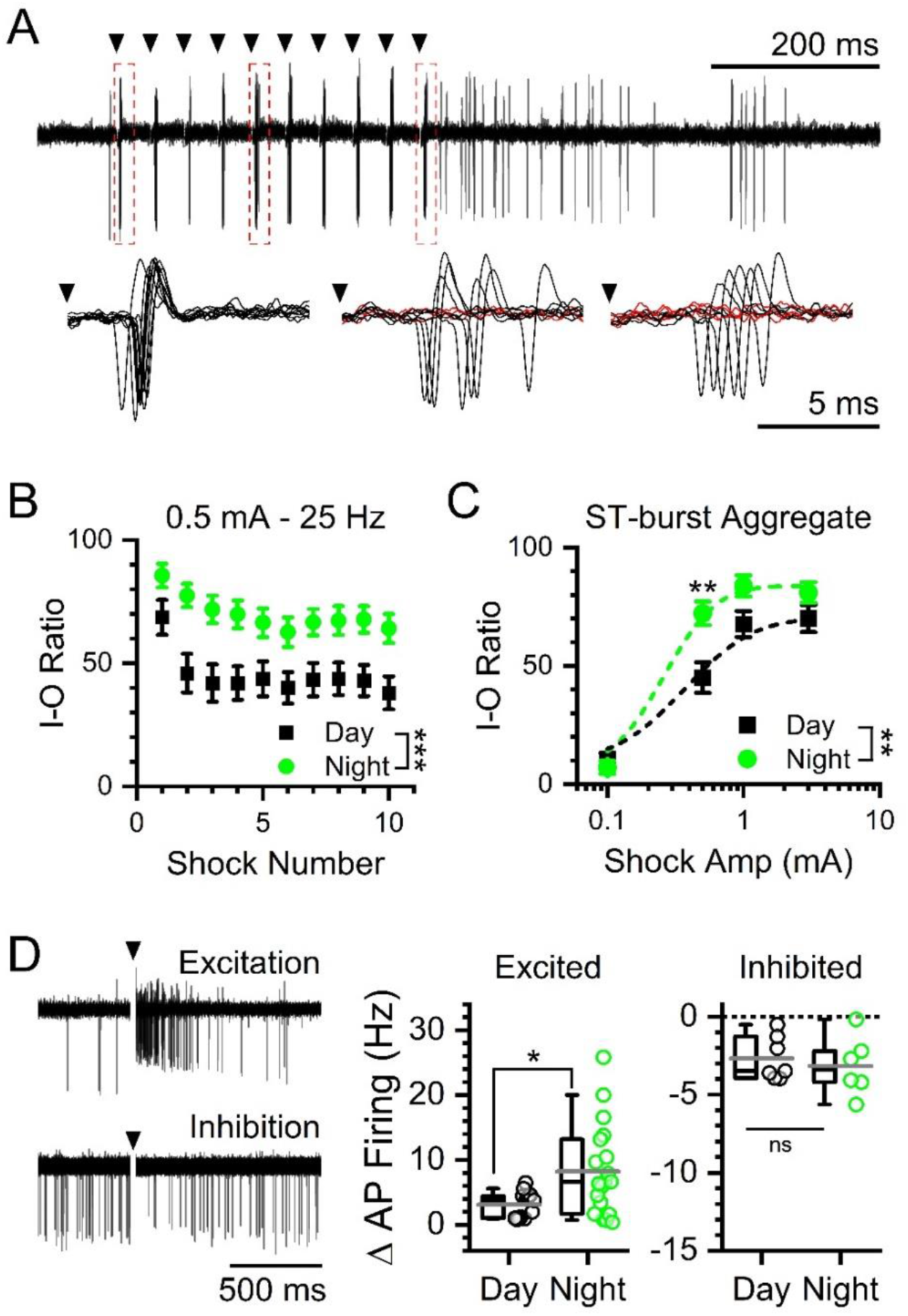
Paradoxical increase in afferent-evoked synaptic throughput at night. **A)** Representative traces showing evoked action potentials by 0.5 mA stimulations at 25 Hz from cell-attached recordings of NTS neurons. Insets show individual action potentials evoked after shock 1 (left), shock 5 (middle), and shock 10 (right). Red traces denote failures. **B)** Plot of the throughput (% successes) of individual stimulations over a train of 10 ST shocks at 0.5 mA stimulation intensity during daytime (ZT5-9) and nighttime (ZT17-21) recordings. At 0.5 mA stim intensity there was a statistical interaction between stimulus number and timed of day with throughput was significantly higher during the night (2-way ANOVA, N = 30 – 39 neurons / 6 mice, ***P < 0.001). **C)** Recruitment curve of action potential throughput at increasing stimulation intensities. There was a significant interaction between time of day and stimulus intensity with throughput was higher at night (2-way ANOVA, N = 30 −39 neurons / 6 mice, **P = 0.008). Post hoc analysis showed a significant difference in throughput at an intermediate stimulus intensity (0.5 mA) (Bonferroni corrected T-Test, N = 30 – 39 neurons / 6 mice, **P = 0.006) but not at very high (1 mA: P = 0.25 and 3 mA: P = 0.72) or low intensities (0.1 mA: P = 0.71). **D)** Representative traces of excited (top) and inhibited (bottom) post-simulation responses. Neurons recorded during the night had a significantly larger post-stimulation excitatory responses compared to day (T-Test, N = 13 – 19 neurons / 6 mice, *P = 0.03). There was no significant difference in the less common inhibitory responses between day or night (T-Test, N = 6 – 7 neurons / 6 mice, P = 0.61).

Immediately following ST synchronized action potentials, many of the NTS neurons showed potentiated firing (44%) with fewer showing transient inhibition (20%) (**Figure 5D, left panel**). Excited neurons had significantly larger post-stimulation action potential burst frequencies during the night compared to neurons recorded in the day (**Figure 5D, middle panel**). While neurons that were inhibited showed no difference in response between time of day (**Figure 5D, right panel**). These data, together with ST synchronized throughput analysis, demonstrate an unexpected greater responsiveness off NTS neurons to afferent driven signals during the night.

### Rhythmic changes in solitary tract (ST) evoked synchronous glutamate release

One potential explanation for increased afferent throughput would be for evoked glutamate release to be regulated differently from spontaneous and enhanced during the night while diminished during the day. Spontaneous and evoked vesicle release pathways are known to have many points of opposing control which may explain the differences in spontaneous versus afferent signaling across day and night (Kavalali et al., 2015). We examined primary afferent synaptic transmission by stimulating the ST and recording synchronous EPSC responses onto second order neurons (**Figure 6**). For each neuron we carefully isolated single afferent inputs using stimulus recruitment protocols; allowing for dissection of release parameters and synaptic performance with minimal contamination from convergent direct and indirect synaptic inputs (Peters et al., 2010; Bailey et al., 2006). Suprathreshold stimulation reliably produced monosynaptic EPSCs which showed no significant difference in EPSC latency to onset (Day: 5.0 ± 0.3 ms vs. Night: 4.8 ± 0.3 ms, T-Test, N = 22 – 23 neurons / 11 mice, P = 0.69) nor synaptic jitter (Day: 105 ± 9 μs vs. Night: 115 ± 9 μs, Mann-Whitney Ranked Sum Test, P = 0.35) between day and night recordings (**Figure 6A**). While evoked EPSC amplitudes varied greatly across neurons (Ranging Day: 72 - 350 pA and Night: 26 – 195 pA) the average initial release (EPSC1) was significantly larger during the day compared to the night (**Figure 6B**). Fluctuation analysis of EPSC1 found an increased covariance (CV) and decreased 1/CV^2^ in recordings during the night consistent with presynaptic changes in either release probability (*Pr*), number of release site (N), or both (**Figure 6C-D**). Trains of stimuli revealed an increased paired-pulse ratio (PPR) between EPSC1 and 2 during the night but no significant differences in steady-state release (**Figure 6E-F**). These data together suggest that across the day afferent-evoked glutamate release undergoes functional presynaptic changes in initial release parameters with no changes in sustained release mediated by vesicle trafficking.

**FIGURE 6:**
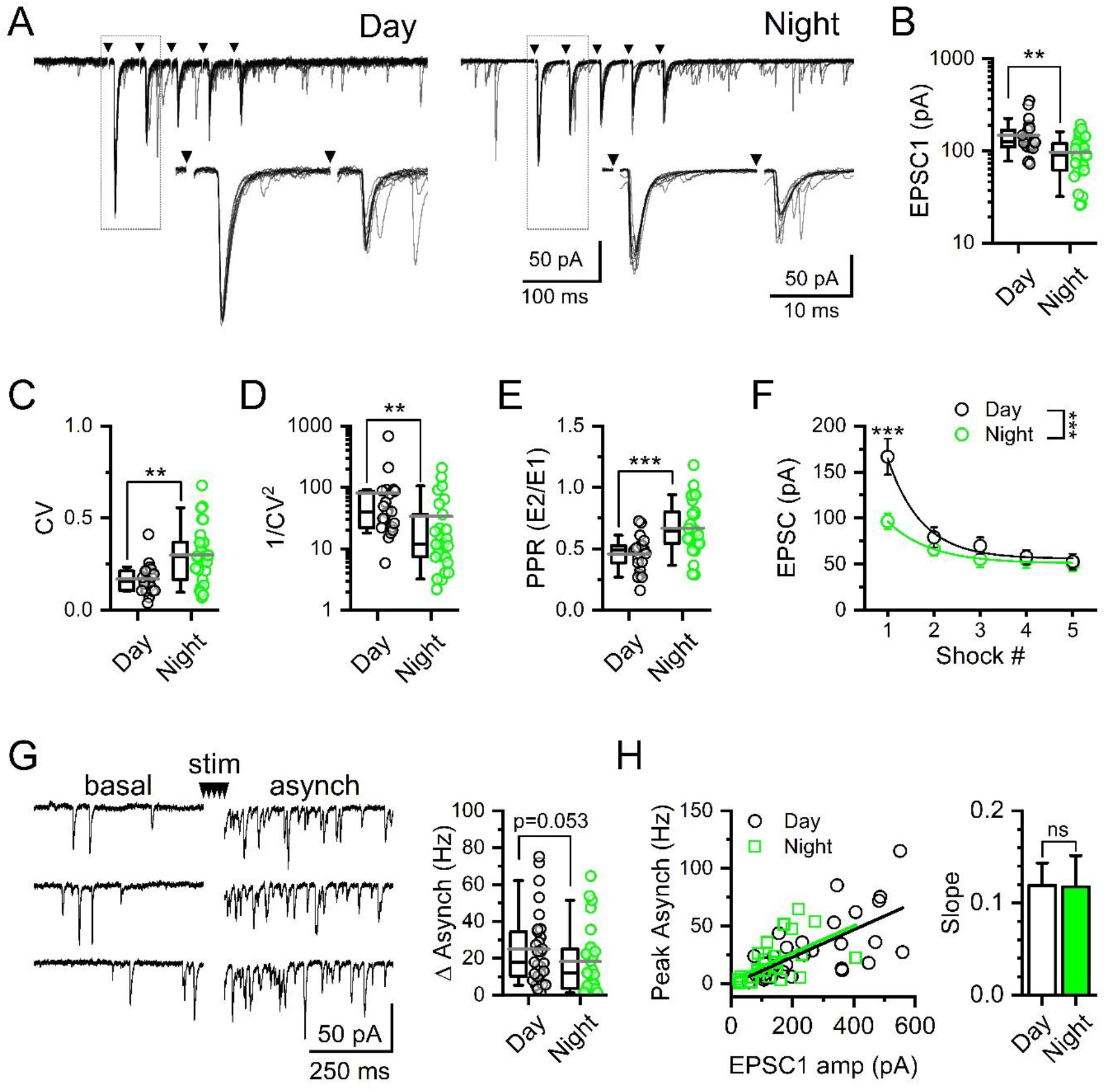
Day / night differences in evoked synchronous glutamate release consistent with presynaptic regulation of the probability of release. **A)** Representative current traces showing 25 Hz suprathreshold stimulation of solitary-tract (ST) afferents and resulting synchronous EPSCs (ST-EPSCs) recorded from slices taken in the day compared to the slices taken at night. Arrows denote time of ST shocks with traces shown as an overlay of 10 individual trials. **B)** The average ST-EPSC amplitude was significantly smaller in recordings made at night compared to the day (T-Test, N = 23 – 27 neurons / 11 mice, **P = 0.002). **C)** The amplitude variability of EPSC1, Coefficient of variance (CV), was significantly higher in recordings made at night (Mann-Whitney Rank Sum Test, **P = 0.004). **D)** Due to this difference in CV, 1/CV^2^ was significantly higher during the day compared to the night (Mann-Whitney Rank Sum Test, **P = 0.004). **E)** As a result of changes in EPSC1, the paired-pulse ratio (PPR) was significantly higher at night compared to the day (T-Test, N = 23 – 27 neurons / 11 mice, ***P = 0.0004). **F)** Average ST-EPSC amplitude across five shocks show a significant difference between day and night (2-way RM ANOVA, N = 42 neurons / 11 mice, ***P < 0.001). Post hoc analysis confirmed EPSC1, but not EPSC2-5, was significantly smaller during the night compared to recordings made during the day (Bonferroni corrected T-Test, EPSC1: ***P < 0.001, EPSC2: P = 0.29, EPSC3: P = 0.21, EPSC4: P = 0.66, EPSC5: P = 0.83). **G)** Afferent-evoked asynchronous EPSCs, elicited by a series of five ST shocks, were not statistically different between day and night recordings (Mann-Whitney Rank Sum Test, N = 27 – 31 neurons / 11 mice, P = 0.053). **H)** Plot of asynchronous release as a function of EPSC1 amplitude. There was no significant difference in the slope of the EPSC1 to asynchronous frequency relationship between day and night (Mann-Whitney Rank Sum Test, N = 27 –31 neurons / 11 mice, P = 0.67). Summary data boxplots show the mean (long gray bar) median (short black bar) with 25-75%, and 10-90% whisker length.

Following bursts of stimuli, the majority of afferents onto NTS neurons exhibit a distinct asynchronous release profile (Peters et al., 2010). Asynchronous release peaks immediately following the final stimulation and was calculated using the first 100 ms bin. We found asynchronous release was lower during the night but just missed statistical significance (**Figure 6G**). The magnitude of asynchronous release scales with size of the synchronous EPSC and was likely diminished due to the decrease in evoked release at night, as there was no change in the slope of the asynch / EPSC1 relationship (**Figure 6H**) (Peters et al., 2010)). Thus, because evoked release was suppressed at night, the asynchronous release profile also tended to be smaller; however, there were not obvious day / night differences specific to this release pathway.

### Diurnal changes in membrane conductance and intrinsic neuronal excitability

In addition to synaptic inputs, neuronal excitability can be modulated intrinsically via changes to membrane conductances (**Figure 7**). Using voltage-clamp, we first measured current responses to depolarizing voltage steps in neurons recorded during the day and night (**Figure 7A**). There was an increase in the resting membrane current (−100 to −60 mV) during the day compared to night as determined using a current-voltage (I-V) plot (**Figure 7B**). From these data we calculated the slope conductance for each neuron and found it was significantly higher during the day (**Figure 7C**). Using current-clamp, we next assayed the ability of injected current to depolarize the neuron and produce action potential firing (**Figure 7D**). NTS neurons showed a subtle but statistically significant increase in action potential firing frequency during the night (**Figure 7E**). These changes in conductance and excitability suggest that intrinsic membrane properties shape neuronal output in addition to synaptic inputs.

**FIGURE 7:**
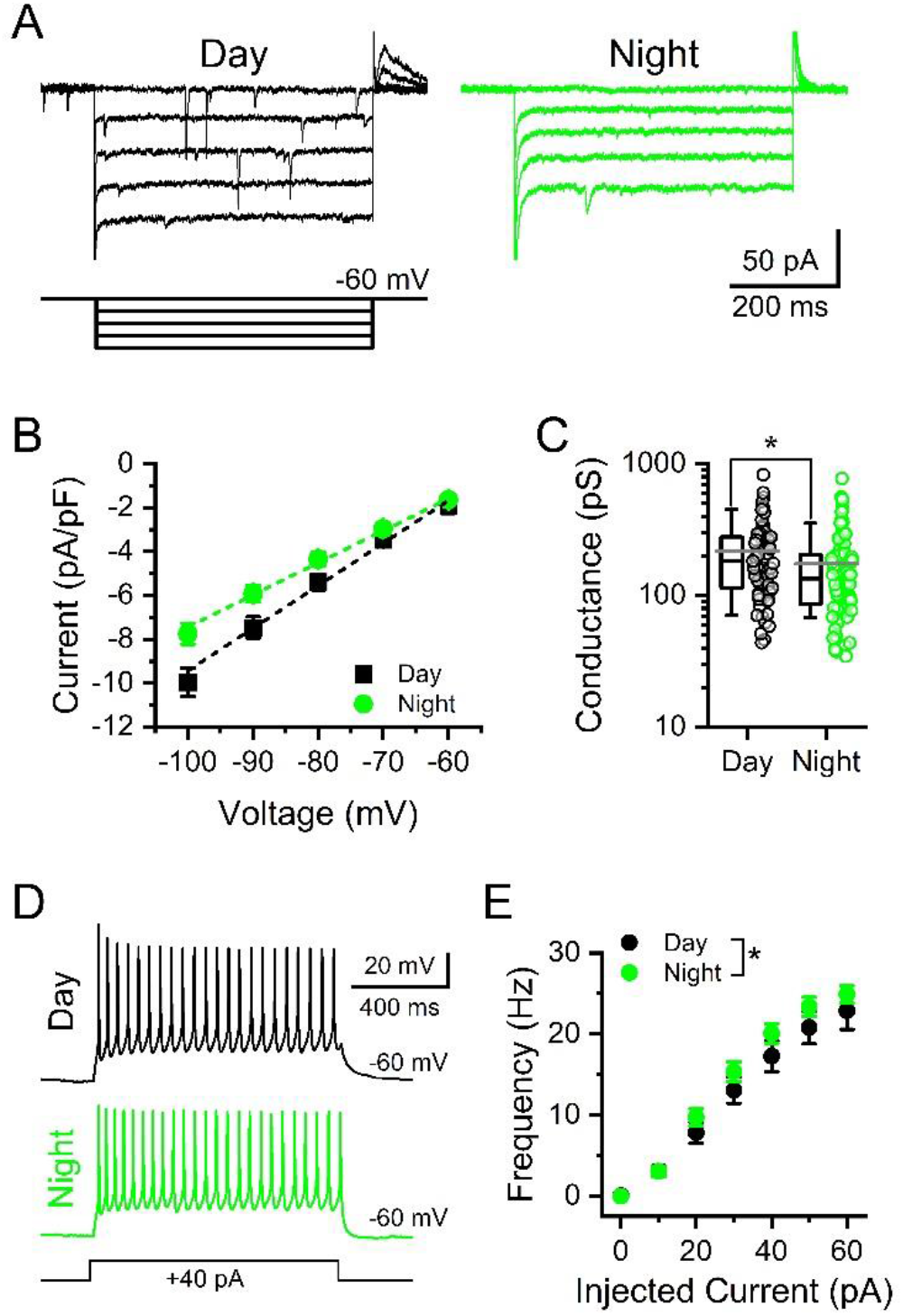
Rhythmic changes in postsynaptic NTS whole-cell conductances impact action potential firing sensitivity to injected current. **A)** Voltage-clamp current traces from NTS neurons demonstrating the current-voltage (I-V) relationship in NTS neurons during day and night. Inset shows the step protocol with 10 mV step intervals from −100 mV to −60 mV. **B)** Plot of the average I-V curves from recordings during the day and night. Values are the average ± stdev of steady-state current density. **C)** The average calculated slope conductance of neurons recorded at night was lower than that of neurons recorded during the day (T-test, N = 67 – 92 neurons / 14 mice, *P = 0.017). **D)** Representative voltage traces from NTS neurons held at −60 mV showing action potential firing elicited during a 1 s, 40 pA, current injection during day (top) and night (bottom) recordings. **E)** Summary plot showing the subtle, but statistically significant, differences between day and night in the action potential firing rate in response to increasing current injections (Mann-Whitney Rank Sum Test, N = 32 – 35 neurons / 6 mice, *P = 0.026).

## DISCUSSION

The role of circadian rhythms in regulating the timing of physiological processes and behavioral responses is established, and the neurophysiological mechanisms mediating these changes remain of considerable interest. Here, we demonstrate that glutamatergic neurotransmission onto NTS neurons is under circadian regulation with peak release during the light phase. This rhythm coupled with daily changes in NTS membrane conductance drives day/night variation in basal and afferent-evoked neuronal firing. We provide evidence for a temporally coordinated shift from high-level constitutive action potential firing during the light to enhanced vagal afferent-evoked synaptic throughput during the dark (Figure 8). These fundamental circadian shifts in synaptic release pathways and synaptic throughput help explain important differences in autonomic regulation that occurs across the day/night cycle and provides a unique mechanism by which circadian rhythms regulate neuronal circuit performance.

**FIGURE 8:**
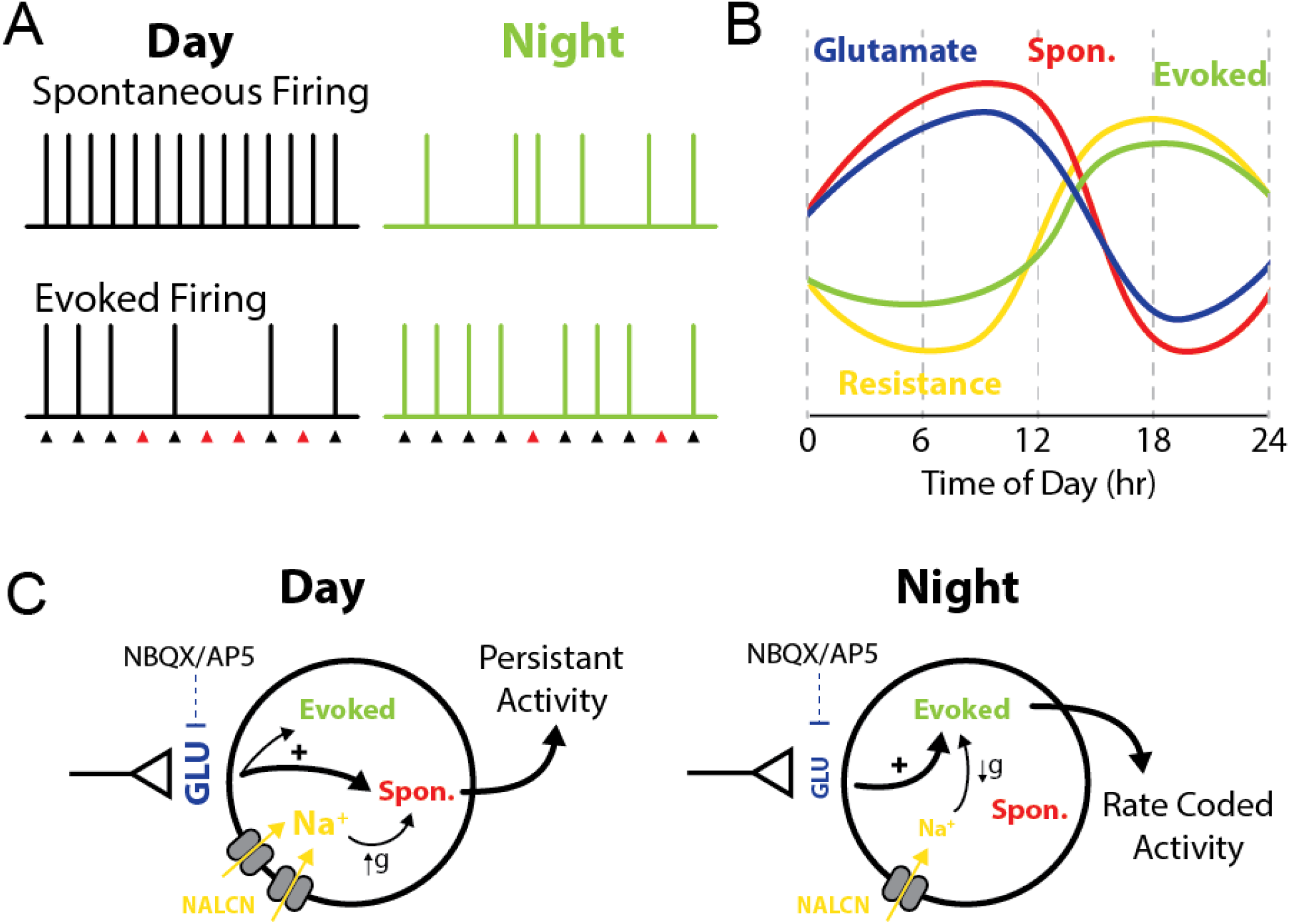
Changes in glutamate release and intrinsic membrane properties shift synaptic processing in the NTS. Model diagram summarizing the observed changes in synaptic function across time of day. **A-B)** During the light phase high spontaneous glutamate release drives elevated postsynaptic action potential firing, while increased postsynaptic membrane conductance diminishes afferent driven throughput. Together this produces a high tone / low responsive synaptic state. In contrast, during the night spontaneous glutamate release is low resulting in decreased basal action potential firing. However, decreased postsynaptic membrane conductance facilitates a high fidelity of synaptic throughput and potentiated post-stimulation bursts. Thus, at night, the synapse has low basal tone and is highly responsive to incoming vagal afferent signals. **C)** Schematic drawing proposing the relationship between each oscillating component within ST afferents and NTS neurons. The axon terminal represents vagal afferents from the solitary tract. Increased glutamate release and membrane conductance enhance firing at rest during the day; while, reduced glutamate release during the night combined with increased membrane resistance enhances afferent evoked AP firing. The symbol ‘g’ represents a change in conductance/membrane resistance.

### Primary visceral afferent and NTS Clock gene expression

In this study we observed a clear rhythm of many clock genes in both the nodose ganglia and the NTS. This confirms studies that have previously identified clock rhythmicity in the hindbrain (Kaneko et al., 2009; Herichova et al., 2007; Chrobok et al., 2020) and nodose ganglia (Kentish et al., 2013) and further corroborates the role of the vagal afferent hindbrain circuit as a circadian oscillator. We demonstrate that *Per1, Per2*, and *Nr1d1* are in phase between NTS and nodose ganglia neurons. Whereas *Bmal1* was only rhythmic in the nodose ganglia and *Clock* was not rhythmic in either tissue. The molecular clock regulates numerous cellular signaling pathways that are involved in the control of cellular excitability. For example, in the SCN calcium rhythmicity is abated in *Cry1/Cry2* DKO or Bmal1 KO mice (Noguchi et al., 2017; Enoki et al., 2017). The increasing evidence that this circuit expresses clock rhythmicity indicates a likelihood for similar cellular rhythmicity to the SCN. Previously in the NTS only the rhythmicity in neuronal action potential firing has been explored (Chrobok et al., 2020). Our current findings on synaptic release and synaptic throughput extend and develop our understanding of the circadian coordination of neurophysiology.

### Circadian regulated spontaneous glutamate release drives tonic NTS activity

Here we show that spontaneous glutamate release onto NTS neurons is increased during the light and diminished during the dark, and that this difference persists in constant conditions and in the presence of TTX. Based on previous studies it is possible that this type of rhythm is synapse specific. In the SCN there is conflicting evidence as to whether synaptic release, both glutamate and GABA, is rhythmic (Itri et al., 2004; Lundkvist et al., 2002; Michel et al., 2002). In another case, glutamate release showed diurnal rhythmicity onto lateral habenula (LHb) neurons but not onto hippocampal neurons (Park et al., 2017). One feature that may contribute to the observed rhythm in the present work is the intrinsically high rates of spontaneous release at solitary tract terminals due to the presence of TRPV1 (Shoudai et al., 2010). TRPV1 provides a unique point of control over this synapse because of its high presynaptic expression and role in mediating quantal release (Peters et al., 2010; Shoudai et al., 2010). It has been shown that TRPV1 mRNA expression is under circadian control suggesting circadian changes in the channel could regulate synaptic release (Kimura et al., 2019; Yang et al., 2015)

Another point of control is calcium, however a role for VGCCs in controlling spontaneous release is still under debate. Some studies in the NTS have determined that spontaneous release is insensitive to VGCC block (Fawley et al., 2016) while others have found VGCCs can regulate spontaneous release (Kline et al., 2019). VGCCs have also been shown to regulate circadian rhythms of firing rates in SCN neurons providing another potential role in regulating neurophysiological rhythms (Enoki et al., 2017b; Pennartz et al., 2002).

Previous studies using organotypic coronal brainstem slices demonstrated a robust circadian rhythm in spontaneous and evoked firing of NTS neurons (Chorbok et al., 2020). The current study demonstrated a rhythm of spontaneous neurotransmission, and that this is sufficient to drive rhythmicity of spontaneous action potential firing of NTS neurons. Spontaneous neurotransmission conveys a large portion of charge transfer in the NTS and selectively increasing or decreasing spontaneous neurotransmission by manipulating temperature modulates action potential firing respectively (Shoudai et al., 2010). This coupling between synaptic transmission and action potential firing has been indicated many times in the SCN. Reducing excitatory postsynaptic activity in the SCN diminishes rhythmicity of SCN firing frequency indicating glutamate may play a role in driving circadian neuronal activity (Lundkvist et al., 2002b). Further, blockade of glutamate receptors reduces both the firing rate and calcium levels of SCN neurons (Michel et al., 2002). This reveals a similarity between the NTS and SCN for a role for glutamate in amplifying circadian firing rates. Another possible role for glutamate is revealed in cultured SCN neurons which only express synchronized circadian firing rhythms once they form synaptic connections (Shirakawa et al., 2000). Whether rhythmic NTS glutamate signaling is a consequence, cause, or both to brainstem circadian rhythmicity remains to be determined.

### Afferent evoked release shows rhythmic changes in release probability

ST-NTS synapses exhibit a high initial release probability (Pr) which is evident by robust frequency dependent depression in response to a train of stimulation (Bailey et al., 2006). Here, we demonstrate diurnal rhythmicity in ST-evoked neurotransmission with peak release occurring during the light. During the dark, we observed a reduced initial evoked EPSC amplitude accompanied by increased EPSC variance and PPR. The coordinated change in both the variance and PPR is indicative of a presynaptic reduction in Pr. Pr can be modulated by many factors and the exact origin of this change is still uncertain. Many proteins that have been shown to regulate presynaptic vesicle fusion, release, and recycling are known to be rhythmic. For example, expression of several voltage gated calcium channels (VGCCs), such as P/Q-type, T-type, and L-type, show circadian rhythmicity in the SCN and cerebellum (Ko et al., 2007; Nahm et al., 2005). In SCN neurons, VGCCs contribute to rhythmic spike generation and intracellular calcium oscillations (Pennartz et al., 2002; Colwell et al., 2000). Other presynaptic proteins, including those that mediate vesicle recycling (Synapsin-I and II), and vesicle fusion (SNAP25, and Munc18), are also rhythmic (Deery et al., 2009; Panda et al., 2002). Alternatively, Pr can be impacted by numerous neuromodulators including corticosterone (Ragozzino et al; 2020), oxytocin (Peters et al., 2008), and vasopressin (Bailey et al., 2006), all of which are under clock control (Kalsbeek et al., 1992; Gillette et al., 1987; Zhang et al., 2011). However, it is unlikely these factors are present at physiological concentrations during *ex vivo* recordings, thus we predict there is likely an *in vivo* role of neuromodulators to amplify or diminish this rhythm. Regardless of the specific mechanism involved, circadian regulation of presynaptic Pr provides a strong leverage point for controlling circuit activity and constitutive firing.

### Intrinsic membrane properties gate neuronal excitability

One of the more striking observations we made is that peak spontaneous and afferent-evoked action potential firing occur out of phase with each other. This provides an interesting contrast to presynaptic release as it reveals a shift in the preferential mode of signaling between day and night. The neurophysiological control for this difference appears to be changes in the postsynaptic Na^+^ leak conductance given our observations of time-of-day changes in the resting membrane conductance of the recorded NTS neurons. Specifically, the membrane conductance is lower during the night producing a ‘tighter’ membrane more responsive to current injections as a function of Ohm’s law. We observed this with the afferent-driven synaptic throughput experiments as well as with directly injecting current and determining the resulting action potential firing frequencies. Previous work has demonstrated that changes in this conductance are is well known to control basal neuronal firing and underly rhythmicity across varying time scales (Impheng et al., 2021; Lu et al., 2007; Simasko et al., 1994). Further, background membrane conductances are proposed to couple and produce an oscillating “bicycle” model where the opposite regulation / expression of resting K+ and Na+ conductances drive rhythmic firing and neural excitability in circadian pacemaker neurons (Flourakis et al., 2015). The extent to which this occurs in NTS neurons is not clear, although we did not measure any obvious changes in K+ conductances. As a consequence of these coordinated presynaptic changes in release probability and postsynaptic changes in membrane conductance, the NTS neurons exhibit this paradoxical increase in intrinsic excitability and synaptic throughput during the dark phase; whereas during the light basal firing is elevated while afferent synaptic throughput is reduced. These coordinated pre-/ postsynaptic changes encode nuanced control over synaptic performance and pair NTS action potential firing and vagal throughput with time of day.

### Functional consequences of rhythmic neurophysiological changes

Many components of the parasympathetic nervous system are under circadian control and previous reports have found that signals that act in the NTS to control metabolic function, including CCK and ghrelin, are rhythmic (Chrobok et al., 2020). CCK effects on food intake also show diurnal variation with peak effectiveness during the inactive phase of the animal (Kraly, 1981). We propose one possible role for rhythmic changes in spontaneous glutamate release and afferent synaptic throughput at this synapse is to provide necessary anticipatory information of rhythmic changes in physiological state including cardiac, respiratory, and digestive rhythms. Increased vagal activity, or parasympathetic tone, is associated with reduced heart rate and respiratory rate, and increased digestive function all of which occur during the inactive phase of the animal (Purnell et al., 2020; Vachon et al., 1987; Hu et al., 2008). This coincides with the observed higher rates of synaptic transmission and basal NTS firing during the light. Furthermore, it is possible that the switch from passive to active signaling enables efficient transmission of viscerosensory signals during the active phase of the animal. Additionally, the NTS projects to the SCN indicating that viscerosensory and interoceptive signals can potentially modulate the clock and that the SCN is likely part of a neural feedback loop which may be involved in regulating autonomic activity (Buijs et al, 2003; Buijs et al., 2014).

## MATERIALS AND METHODS

### Animals

Adult male C57BL/6N mice (20 - 30 g) were obtained from either Envigo or Jackson laboratories. Mice were maintained under standard 12:12 hour light – dark (LD) cycle in a temperature-controlled (23 ± 1 °C) room with *ad libitum* access to water and standard pellet chow. For experiments involving constant darkness (12:12 hour dark – dark (DD), animals were house in identical conditions except for constant darkness for 24-42 hours prior to experimentation. All experiments were performed in accordance with procedures approved by the Institutional Animal Use and Care Committee (IACUC) at Washington State University.

### Molecular biology

#### Tissue preparation

Mice were deeply anesthetized and euthanized at six time points across the circadian cycle: ZT0, ZT4, ZT8, ZT12, ZT 16, and ZT 20 (N = 4 mice / time-point, 24 mice total). Brainstems and nodose ganglia were isolated and flash frozen on dry ice and stored at −80°C. Brainstems, separated rostral to the cerebellum, were mounted on a Leica cryostat and cut on the coronal plane producing 3 x 300 μm thick slices that were direct mounted onto frozen glass slides. From these slices we collected NTS enriched tissue using a 0.5 mm diameter tissue punch. All collected tissues were stored in RNAlater (ThermoFisher).

#### RT-qPCR

The tissue was processed for RNA extraction using Qiazol extraction and RNeasy Micro Kit (QIAGEN). Concentrations of mRNA were determined using spectrophotometry with samples diluted to the same final concentration. Immediately following extraction, mRNA was treated with Ambion DNase treatment and removal (Life Technologies) and cDNA synthesis was performed with QuantiTect^®^ cDNA reverse transcription kit (QIAGEN). Reverse-transcription qPCR assays were performed using TaqMan chemistry commercially available validated assays from Life Technologies. Transcripts were amplified on a 7500 Fast Real-Time PCR System by Applied Biosystems (Life Technologies) using TaqMan probes for *Rn18s* (control gene; Mm03928990_g1), *Per1* (Mm00501813_m1), *Per2* (Mm00478099_m1), *Bmal1 (Arntl1*) (Mm00500226_m1), *Nr1d1* (Rev-erbα) (Mm00520708_m1), *Clock* (Mm00455950_m1), *Cckar* (Mm00438060_m1), *Cckbr* (Mm00432329_m1). Samples were run in triplicate with a standardized 5 ng (2.5 ng / μL x 2 μL) of cDNA and compared using the ^ΔΔ^CT method of relative quantification, with *Rn18s* used as the control housekeeping gene and ZT0 values as the relative target. Kruskal-Wallis test with post-hoc multiple comparison was performed for statistical analysis.

### Slice electrophysiology

#### Horizontal brainstem slice preparation

Brainstem slices were isolated from mice deeply anesthetized with isoflurane as previously described (Appleyard et al., 2005). The brainstem was removed from just rostral to the cerebellum to the first cervical vertebrae and placed in ice-cold artificial cerebral spinal fluid (aCSF) containing (mM): 125 NaCl, 3 KCl, 1.2 KH_2_PO_4_, 1.2 MgSO_4_, 25 NaHCO_3_, 2 CaCl_2_, and 10 dextrose, bubbled with 95% O_2_ – 5% CO_2_. aCSF was brought to a pH of 7.40 using 1M HCl. Once chilled, the tissue was cut to remove the cerebellum and the tissue block was mounted horizontally to a pedestal with cyanoacrylate glue and submerged in cold aCSF on a vibrating microtome (Leica VT1200S). Approximately 150-200 μm was removed from the dorsal surface and then a single 250 μm thick horizontal slice was collected containing the solitary tract (ST) along with the neuronal cell bodies of the medial NTS region. Slices were cut with a sapphire knife (Delaware Diamond Knives, Wilmington, DE) and secured using a fine polyethylene mesh in a perfusion chamber with continuous perfusion of aCSF bubbled with 95% O_2_ – 5% CO_2_ at 32 °C. Brainstem slices were generated at 6-hour intervals, starting at zeitgeber time (ZT) 0, and data were binned with reference to the time of each recording; either every 4 hours or into day (ZT0-12) and night (ZT12-24). In some experiments where only two time points were used, slices were taken between ZT4-5 and ZT16-17.

#### Whole cell patch-clamp recordings

Recordings were performed on NTS neurons contained in horizontal brainstem slices using an upright Nikon FN1 microscope with a Nikon DS-Qi1Mc digital camera and NIS-elements AR imaging software. Recording electrodes (3.0 – 4.5 MΩ) were filled with one of two intracellular solutions containing either (Cs-internal, mM): 10 CsCl, 130 Cs-Methanesulfonate, 11 EGTA, 1 CaCl_2_, 2 MgCl_2_, 10 HEPES, 2 Na_2_ATP, and 0.2 Na_2_GTP; or (K+ internal, mM) 6 NaCl, 4 NaOH, 130 K- gluconate, 11 EGTA, 1 CaCl_2_, 1 MgCl_2_, 10 HEPES, 2 Na_2_ATP, and 0.2 Na_2_GTP. For measurements of excitatory post-synaptic currents (EPSCs), neurons were studied under voltage-clamp conditions with an MultiClamp 700A amplifier (Molecular Devices, Union City, CA) and held at V_H_ = −60 mV in whole-cell patch configuration. Only recordings with a series resistance of <20 MΩ were used for experiments to ensure good access and maintenance of voltage-clamp. Signals were filtered with a 1 kHz bezel filter and sampled at 20 kHz using Axon pClamp10 software (Molecular Devices).

#### Extracellular recordings

To monitor action potential firing in single NTS neurons with unperturbed intracellular conditions we utilized loose-seal extracellular recordings. Recording pipets (3.0 – 4.5 MΩ) were filled with the standard extracellular aCSF and a relatively low resistance (loose) seal was formed with the NTS neuron. The typical seal resistance was between 10 – 20 MΩ and neurons were recorded in the current-clamp configuration with no injected current. This allowed for stable recordings and the clear resolution of single-unit extracellular spikes resulting from spontaneous and evoked action potential firing.

#### Solitary tract stimulation

Second-order neurons in the NTS receive direct innervation from ST afferents, which include the primary vagal afferent terminals, and can be identified based on the precision of shock evoked synchronous glutamate release. To selectively activate ST afferent fibers, a concentric bipolar stimulating electrode (200 μm outer tip diameter; Frederick Haer Co., Bowdoinham, ME, USA) was placed on distal portions of the visible ST rostral to the recording region. We delivered a train of five current shocks to the ST (60 μs shock duration at 25 Hz, 6 s inter-stimulus interval) using a Master-8 isolated stimulator (A.M.P.I., Jerusalem, Israel). Input latency, the time between the shock artifact and onset of the synchronous EPSC, and synaptic “jitter”, the standard deviation of the latency, were measured to identify monosynaptic inputs. Jitters of <200 μs identify monosynaptic afferent inputs onto the NTS neuron (Doyle and Andresen, 2001).

#### Data analysis

For brainstem slice recordings the digitized waveforms of action potentials and synaptic events were analyzed using an event detection and analysis program (MiniAnalysis, Synaptosoft, Decatur, GA). Analysis for all ST-stimulated currents was accomplished using Clampfit 10 (Molecular Devices). For quantal, spontaneous glutamate release, all events >10 pA were counted for frequency values. Fitting of quantal EPSC amplitudes and decay kinetics (90-10%) was performed using a fitting protocol (MiniAnalysis) on > 500 discrete events or 5 minutes of recording time if less than 500 events occurred.

#### Chemicals and drugs

Drugs were purchased from retail distributers including tetrodotoxin (TTX, 1 μM; Tocris #1069), 2,3-dioxo-6-nitro-7-sulfamoyl-benzoquinoxaline (NBQX, 25 μM; Tocris #0373), D-2-Amino-5-phosphonopentanoic acid (D-AP5, 25 μM; Tocris #0106). These were stored as stock aliquots and diluted in fresh aCSF to their final concentration the day of recording. The general salts used for making bath solutions were purchased from Sigma-Aldrich. All drugs were delivered via bath perfusion at a flow rate of ~2 mL / min and slices were allowed to fully recover before the start of recording.

### Statistical analyses

For statistical comparisons we used Sigma Stat software (Systat Software Inc., San Jose, CA). The data were tested for normality and equal variance and the appropriate parametric or nonparametric statistics were used; including, ANOVA with post hoc analysis, T-tests, Mann Whitney Rank Sum test, and linear regression analysis. Data sets were analyzed using Grubb’s test and extreme outlier values were excluded from statistical comparisons. Specific test used are indicated in the results section. Comparisons were considered statistically different with an alpha level of P < 0.05.

## ACKNOWLEDGMENTS

This work was supported by grants from the National Institutes of Health (DK092651 to JHP) and (DK119811 to INK), and the National Science Foundation (CAREER 1553067 to INK).

## Author contributions

Design experiments (*FJR, INK, JHP*).

Conduct experiments (*FJR, BP, JHP*).

Analyze data (*FJR, BP, INK, JHP*).

Prepare and write manuscript (*FJR, INK, JHP*).

